# Interference Between Motor Memories Arises From Implicit Recalibration

**DOI:** 10.64898/2026.04.23.720436

**Authors:** Aarohi Pathak, Adarsh Kumar, Timothy N. Welsh, Pratik K. Mutha

## Abstract

Interference between consecutively acquired motor memories is a defining feature of sensorimotor adaptation, yet its mechanistic origin remains unresolved. Because adaptation is supported by separable explicit strategies and implicit recalibration, it offers a means to identify the learning process that gives rise to interference. Across four visuomotor adaptation experiments, we examined the conditions under which the acquisition of a new, competing motor memory influences the expression of a previously acquired memory. We selectively biased new learning towards either explicit or implicit processes, and quantified its impact on the recall of the original memory 24-hours later. Under standard adaptation conditions, participants exhibited classic interference, such that re-learning was indistinguishable from naïve performance. However, when new learning was driven primarily by explicit strategies induced through delayed endpoint feedback, interference was markedly attenuated and the original memory was preserved. In contrast, when the competing memory was implicitly forged under error-clamp conditions, robust interference emerged. Furthermore, disrupting posterior parietal cortex (PPC) with cathodal hd-tDCS prior to implicit learning attenuated interference, indicating that intact PPC processing is required for incorporating new learning into an existing sensorimotor representation. Taken together, these findings suggest that interference reflects the integration of new learning into a shared representational substrate via implicit recalibration, a process that limits the coexistence of competing motor memories.

## INTRODUCTION

Interference, the diminished ability to acquire or retrieve information due to competing experiences, is a fundamental constraint across multiple memory systems. In verbal memory, for example, learning a new list of words can impair recall of a previously learned list, and past learning can hinder, in an anterograde manner, the encoding of new information (Keppel & Underwood, 1962; Postman, 1962; Underwood, 1957). Similarly, in perceptual learning, training on one visual discrimination task can disrupt improvements on a closely-related task (Seitz et al., 2005; Shibata et al., 2017; Yotsumoto et al., 2009). Even in higher-level domains such as problem solving or language acquisition, previously established rules or associations can bias performance and block the integration of new information (Schneider & Simonsmeier, 2025; VanDyke, 2007). Interference is shaped by factors such as representational similarity, task context, and temporal spacing between learning episodes, and it is believed to be the behavioural outcome of an underlying competition for shared neural or cognitive resources (Bock et al., 2005; Bouton, 1993; Kumar et al., 2018; Wada et al., 2003).

Interference is also a defining constraint within the motor system. Motor learning leads to the formation of motor memories, often through the recalibration of existing sensorimotor mappings, and interference is known to occur when these mappings conflict. For example, studies of motor adaptation, a form of motor learning driven by error, have revealed that the memory acquired by adapting to a novel force field can be disrupted by subsequent exposure to an opposing force field (Brashers-Krug et al., 1996; Karniel & Mussa-Ivaldi, 2002; Stockinger et al., 2014). Likewise, adaptation to a new visuomotor rotation can hinder the retention of an opposite rotation learned earlier (Caithness et al., 2004; Hinder et al., 2007; Krakauer et al., 2005; Miall et al., 2004; Wigmore et al., 2002). Collectively, these findings point to interference as a robust feature of adaptation, but leave unresolved the processes through which competing motor memories interact.

Unlike some cognitive domains where the underlying mechanisms are hard to discern, motor adaptation is known to be driven by at least two separable processes, the contributions of which can be independently assessed (Mazzoni & Krakauer, 2006; Oza et al., 2024; Taylor et al., 2014). This dissociability in the processes provides a unique opening to probe not just whether interference occurs as a phenomenon, but which specific learning process drives it. Motor adaptation emerges from the combined operation of explicit strategies and implicit recalibration processes. Strategies are verbally sensitive, volitional responses which can involve, for example, deliberate re-aiming of the hand away from a target to compensate for an error introduced by a visuomotor rotation (Anguera et al., 2010; Christou et al., 2016; Fernandez-Ruiz et al., 2011; Taylor et al., 2014). In contrast, implicit processes drive automatic, gradual updates to motor plans in response to sensory prediction errors (Bond & Taylor, 2015; Mazzoni & Krakauer, 2006; Morehead et al., 2015; Oza et al., 2024; Tsay et al., 2022; Tseng et al., 2007). These processes differ in terms of timescale of operation (McDougle et al., 2015; Smith et al., 2006), flexibility (Bond & Taylor, 2015; Morehead et al., 2017; Wilterson & Taylor, 2021) and underlying neural substrates (Clower et al., 1996; Mutha et al., 2011; Tseng et al., 2007), and also give rise to distinct phenomena. Strategies are fast-acting, flexible, and depend on working memory (Anguera et al., 2010; Christou et al., 2016; McDougle et al., 2015), while implicit processes are slower, less flexible (Bond & Taylor, 2015; Wilterson & Taylor, 2021) and require intact cerebellar (Tseng et al., 2007) and posterior parietal processing (Kumar et al., 2020; Mutha et al., 2011). Strategy use gives rise to savings or faster re-learning (Haith et al., 2015; Morehead et al., 2015), while implicit recalibration drives interlimb generalization of learning (Kumar et al., 2020; Kumar & Mutha, 2023).

This separability of processes has enabled some studies to probe whether interference preferentially disrupts explicit or implicit components of adaptation (Avraham et al., 2021; Lerner et al., 2020). However, even when interference is decomposed in this way, the emphasis has largely been on identifying which memory expression is altered, rather than on determining which learning process actively generates interference when a competing task is acquired. Here, we address this more fundamental issue of *origin*, asking which of these processes is a more potent source of interference when engaged during new learning. One possibility is that, like savings, interference arises primarily from explicit strategy use. For instance, a strategy used to counter a visuomotor rotation may persist and bias performance in a subsequent, opposing task (Day et al., 2016; Lerner et al., 2020; Wang & Sainburg, 2003). Alternatively, the origin may lie in implicit recalibration, wherein a new, inflexible sensorimotor mapping conflicts directly with an established one, generating interference at the level of implicitly updated motor representations (Avraham & Ivry, 2025). A third possibility is that both processes contribute jointly, with their combined influence producing the observed interference.

In the present work, we first aimed to disentangle these possibilities at the behavioral level. By selectively driving new learning in a visuomotor adaptation task through either explicit or implicit processes, and then quantifying its impact on a previously acquired motor memory, we sought to determine which process serves as the primary source of interference. Our results point to a clear dissociation: robust interference is observed when new learning is dominated by implicit recalibration, but not when it is primarily explicit strategy based. Next, we proceeded to identify the neural locus of interference. Implicit recalibration is known to depend on cerebellar computations of sensory prediction error (Galea et al., 2011; Martin et al., 1996; Tseng et al., 2007), but evidence points to the posterior parietal cortex (PPC) as a key cortical site where the updated mappings are likely integrated and represented (Kumar et al., 2020; Mutha et al., 2011; Tanaka et al., 2009). Imaging studies have shown adaptation-related changes in PPC (Clower et al., 1996; Danckert et al., 2008; Diedrichsen et al., 2005), and neurostimulation work has shown that disrupting PPC processing selectively attenuates implicit learning (Kumar et al., 2020). These findings suggest that if implicit recalibration drives interference as revealed by our behavioral work, and PPC is a key locus for this recalibration, then disrupting the PPC prior to the acquisition of the new memory should reduce the degree of interference. We tested this hypothesis by applying cathodal high-definition transcranial direct current stimulation (hd-tDCS) over the left PPC, and found this to be the case.

Taken together, these findings advance our understanding of interference on two levels. At the behavioral level, they reveal that implicit recalibration, and not explicit strategy use, is the primary driver of interference. At the neural level, the findings implicate the PPC as a key contributor to interference arising from implicit recalibration. Beyond their theoretical significance, these insights hold practical implications for skill acquisition and rehabilitation, where reducing interference may significantly improve training and recovery outcomes.

## METHODS

### Participants

A total of 90 right-handed individuals (56 males, 34 females), aged 18-34 years (mean = 21 years, SD = 2.3), participated in experiments 1-3 (30 participants per experiment). Sample size was informed by prior experiments from our laboratory using similar visuomotor adaptation paradigms and statistical comparisons between rotation conditions (Kumar et al., 2018). Sample size calculations conducted using G*Power (version 3.1.9.6) indicated that a minimum of 7 participants per group would provide 80% power to detect large effects (Cohen’s *d* = 1.4) at an alpha level of 0.05. We deliberately over-recruited relative to this minimum to allow for tests of interactions, effector-related effects, and individual-difference analyses. Half the subjects in each experiment (n=15) were randomly assigned to perform the task with their left arm, while the remaining (n=15) were assigned to perform the task with their right arm. During initial analyses, no main effect or interaction involving arm was observed, and therefore, data were pooled across participants for each experiment.

A separate group of 20 participants was recruited for experiment 4 that employed the hd-tDCS. One participant chose to discontinue stimulation before completing the session and was excluded, resulting in a final sample of 19 participants (12 males, 7 females; mean age = 21 years, SD = 1.4). All participants performed the task using only their right arm because experiments 1-3 revealed no effects of arm used on adaptation or interference.

All participants had normal or corrected-to-normal vision and reported no neurological disorders, cognitive impairment, or orthopaedic injuries. They were naïve to the task paradigm and the overall purpose of the experiments. Handedness was assessed using the Edinburgh Handedness Inventory (Oldfield, 1971). Participants provided informed consent and received financial compensation for their participation. The study procedures were approved by the Institute Ethics Committee at the Indian Institute of Technology Gandhinagar.

### Apparatus

The experimental setup consisted of a HDTV mounted horizontally above a digitizing tablet (Calcomp Inc.). A semi-transparent mirror was positioned between the tablet and the display, allowing visual feedback of the cursor and targets on the mirror while occluding direct vision of the participants’ hand. Participants used a hand-held stylus to perform reaching movements on the tablet, with stylus (hand) position sampled at 120 Hz. Cursor motion was either veridical or rotated relative to hand motion, depending on the task phase. In Experiment 4, hd-tDCS (Soterix Medical) was administered as participants performed the task in this apparatus; stimulation parameters are described later.

### Task Procedure

Participants performed center-out reaching movements from a central start circle (2 cm diameter) to one of eight peripheral targets (2 cm diameter) positioned radially at 45° intervals on a virtual circle with a radius of 15 cm. At the start of each trial, participants moved the cursor to the central start position and held it there for 500 ms. One target then appeared, accompanied by an auditory tone signalling that participants should begin their movement. Participants were instructed to make fast and accurate reaching movements using primarily the shoulder and elbow, avoiding wrist or finger movements. Target order was pseudorandomized such that each of the eight targets appeared once per set of eight trials. This sequence was identical across participants, rotations, and experiments. After each reach, a second auditory tone prompted participants to return to the start position, and a brief numerical feedback (10 points for landing within the target, 5 points for landing within 2.5 cm of the target, and no points for missing the target beyond this range) was displayed for 500 ms. This feedback was included solely to maintain participant engagement and did not influence our analyses or lead to a financial reward for participants.

All experiments followed a “Baseline-A-B-A” structure across two consecutive days. On Day 1, participants first completed a set of 48 baseline trials in which the cursor and hand motion were matched (i.e., veridical visual feedback). Subsequently, they were exposed to a 30° clockwise (CW) visuomotor rotation “A” for 200 trials (A_1_), followed by 200 trials of exposure to the opposite, 30° counterclockwise (CCW) rotation B. No washout trials were interposed between A_1_ and B, ensuring that B learning occurred on top of the adapted state from A_1_. 24-hours after adapting to B (i.e., on Day 2), participants returned to perform 200 trials while experiencing the CW rotation A again (A_2_). For A_1_ and A_2_, all perturbations were abruptly introduced (i.e. the full 30° rotation was applied on the first trial of each rotation block) and cursor feedback was available throughout the movement, whereas the conditions under which B was performed varied across the different experiments. During A_1_ learning, participants were instructed to make the cursor hit the target, while instructions for B depended on the specific experiment (see below). When participants returned for A_2_ on Day 2, they were again instructed to try and make the cursor hit the target.

### Experiment-specific manipulations for rotation B

Our experiments differed in the way in which rotation B was manipulated across experiments.

#### Experiment 1

This experiment was designed to replicate classic demonstrations of interference following exposure to opposing perturbations. Therefore, rotation B consisted of an abruptly introduced 30° CCW rotation with continuous online cursor feedback after 30° CW A_1_ learning. Participants were told to make the cursor hit the target.

#### Experiment 2

In this experiment, rotation B was designed to emphasize explicit strategy use. Cursor feedback during the reach from the start position to the target was removed, and the cursor (rotated 30° CCW) was displayed only as an endpoint, after a delay of 2.5s following movement completion. As in experiment 1, participants were instructed to make this (endpoint) cursor hit the target. By removing continuous visual feedback and providing only endpoint information, participants were encouraged to rely primarily on deliberate, strategic adjustments to counter the perturbation (Dawidowicz et al., 2022).

#### Experiment 3

In experiment 3, rotation B was implemented as an “error-clamp”, wherein the direction of cursor motion was “clamped” to a fixed 30° CCW angle relative to the target direction. In other words, the direction of cursor motion remained invariant to the participants’ actual hand direction, and always followed a fixed trajectory. Participants were explicitly instructed to ignore the cursor and aim their hand directly toward the target (Morehead et al., 2017). Such a manipulation is known to drive implicit recalibration while minimizing the contribution of explicit strategies.

#### Experiment 4

Experiment 4 followed the same error-clamp procedure as in experiment 3 for B learning. In addition, cathodal hd-tDCS was applied over the left PPC. The PPC was selected given that it has been causally implicated in visuomotor learning, particularly adaptation driven by the slowly evolving implicit process (Kumar et al., 2020). Stimulation was delivered using a 4×1 ring electrode configuration centered over P3, with the P3 location derived based on the international 10-20 system (Kuo et al., 2013). During stimulation, a constant current of 2 mA was applied for 20 minutes. Participants remained seated quietly for the first 15 minutes of tDCS delivery, after which stimulation overlapped with the initial phase of B learning (∼50 trials). There was no interruption in the remaining trials of the B block, i.e. participants continued to perform these adaptation trials as the stimulation ended.

### Data Reduction and Analysis

Hand position (stylus) data were low-pass filtered using a Butterworth filter (10 Hz cutoff), and velocity values were obtained by numerically differentiating the position data. Movement onset was identified as the first point in the velocity profile where the velocity fell below 5% of the peak, working backwards from the point of peak velocity towards the start of the trial. Likewise, movement offset was defined as the first point where the velocity fell below 5% of the peak working forwards from the point of peak velocity towards the end of the trial. Additionally, reaction time (RT) was calculated as the interval between target appearance and movement onset, and movement time (MT) as the interval between movement onset and offset.

We also computed the hand deviation at the point of peak velocity for the baseline and adaptation trials. Hand deviation was defined as the angular difference between the vector from the start position to the target and the vector from the start position to the hand position at peak velocity. These deviations were expected to be very small for baseline trials. Baseline hand angles exceeding ±10° were excluded as outliers. Baseline-corrected hand angles were then computed by subtracting the mean baseline hand angle from all subsequent trials. Additional trials were excluded if movement was not initiated, if stylus contact was lost, or if hand angles exceeded ±100°. Across all experiments, less than 0.05% of trials were excluded.

Adaptation was quantified as a change in hand deviation over the adaptation block. For analysis purposes, hand deviation data were binned (1 bin = 8 trials), resulting in 6 baseline bins and 25 bins for each of the adaptation blocks. Early learning was defined as the mean hand deviation on the first bin of the adaptation block, while late learning was defined as the mean hand deviation on the last bin. RT, MT and peak velocity analyses focused on average values computed for an entire block rather than just the early and late phases. In addition to group-level analyses, correlational analyses were performed to assess relationships between individual differences in learning across experimental phases.

Our main statistical analysis focused on the differences in hand deviation between the early and late adaptation phases of the A_1_ and A_2_ blocks. This was done via a two-way repeated measures ANOVA with Block (A_1_, A_2_) and Learning Phase (early, late) as factors. This analysis revealed the effects of the intervening B learning on the prior memory of A. Interference was operationally defined as the absence of an increase in early A_2_ hand deviation relative to early A_1_. Conversely, retention of prior learning was defined as a significant increase in early A_2_ hand deviation compared to early A_1_. Note that the hand deviation reflects the angular deviation of the reach in the compensatory direction. Thus, larger deviations correspond to smaller directional errors (i.e., cursor motion closer to the target direction). Comparisons between RT, MT and peak velocity were done using paired t-tests. All of these analyses were conducted separately for each experiment. Significant main effects or interactions were followed up with Bonferroni-corrected post-hoc comparisons. The significance threshold was set at 0.05.

## RESULTS

### Replication of the classic interference effect

In our first experiment (Figure 1A), participants adapted to a 30° CW rotation (A_1_), followed by adaptation to a 30° CCW rotation (B). After a 24-hour delay, they were retested on rotation A (A_2_). Behavioural patterns during learning were characteristic of classic visuomotor adaptation. On initial exposure to A, cursor trajectories were curved (Figure 1B, red), but progressively straightened as learning proceeded (Figure 1C, red), indicating successful adaptation to the perturbation. Critically, when participants were retested on A after 24 hours (A_2_), their initial trajectories closely resembled those from the early phase of A_1_, suggesting no benefit from prior A learning (Figure 1B, blue). That is, participants exhibited performance consistent with naïve-like learning, with adaptation progressing in a manner similar to A_1_. This trend was also evident in the group-averaged hand deviations (Figure 1D), which showed an increase during A_1_ as subjects adapted to the rotation, then a drift in the opposite direction upon exposure to B, and then a pattern that mirrored A_1_ during the A_2_ retest (Figure 1E).

**Figure 1.**
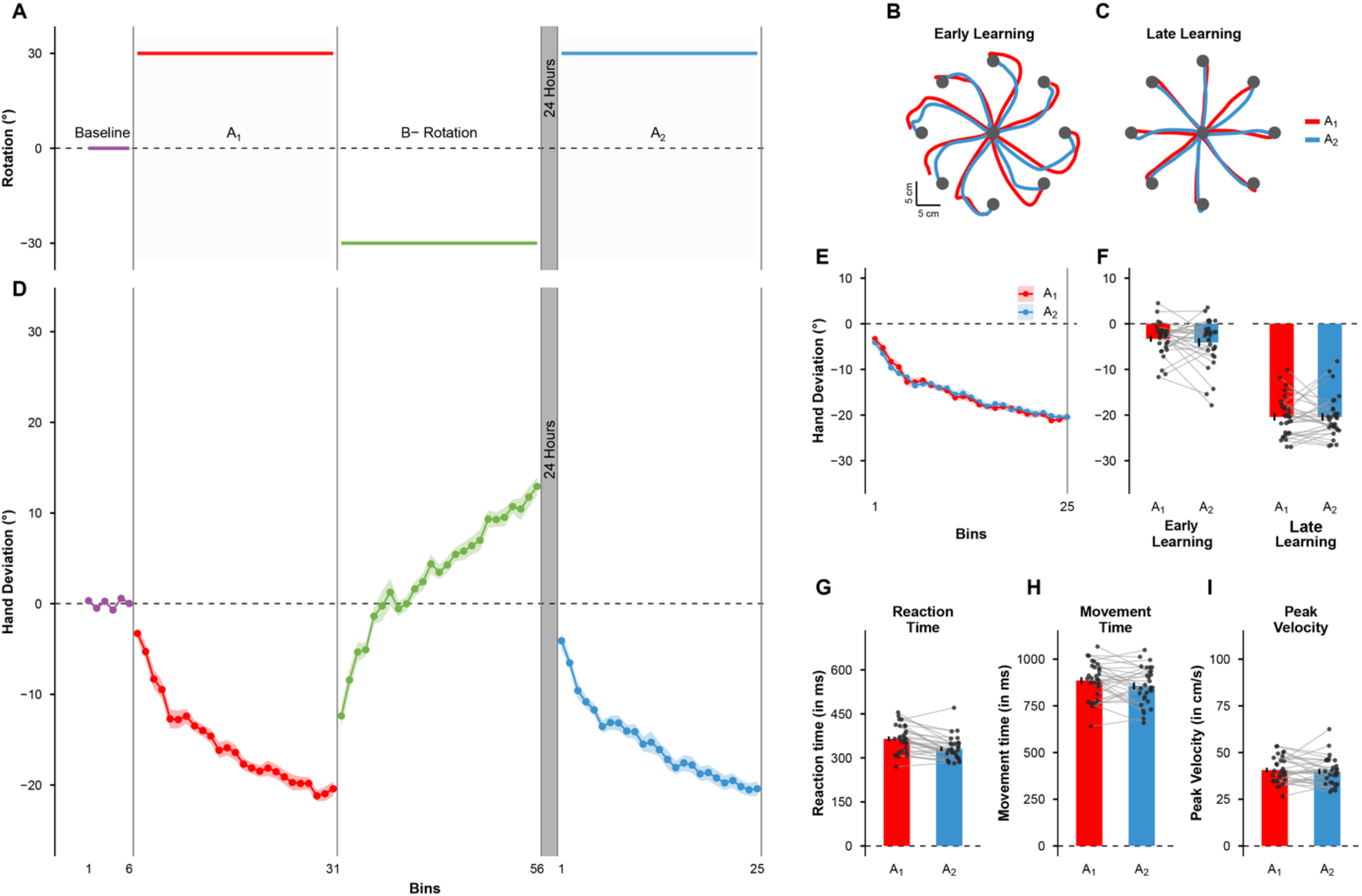
Replication of the classic interference effect. **(A)** Progression of the Baseline-A-B-A task structure. Subjects first perform baseline movements and are then exposed to a 30° visuomotor rotation A (A_1_), followed by exposure to an opposite rotation B, and are then finally retested on A after 24 hours (A_2_). **(B)** Example cursor trajectories from a representative participant during the early phase of learning in A_1_ (red) and A_2_ (blue). **(C)** Example cursor trajectories from the same participant during the late phase of learning in A_1_ (red) and A_2_ (blue). **(D)** Group-averaged hand deviation across bins, showing the Baseline, A_1_, B, and A_2_ blocks (baseline: bins 1-6; A_1_: bins 7-31; B: bins 32-56; A_2_: bins 1-25 following the 24-hour delay). Shaded bands are SEM. **(E)** Group-averaged hand deviation curves for A_1_(red) and A_2_(blue) overlaid together. **(F)** Mean hand deviation during the early (first bin) and late (last bin) phases of A_1_ and A_2_. **(G)** Mean reaction time, **(H)** movement time, and **(I)** peak velocity during A_1_ and A_2_. Dots represent individual participants; bars indicate group means. Error bars represent SEM.

To quantify and test these observations, we conducted a Block (A_1_, A_2_) x Learning Phase (early, late) repeated-measures ANOVA on the hand deviations. Our analysis revealed a significant main effect of Learning Phase (*F*(1,29) = 317.22, *p* < 0.001, *η*_*p*_^*2*^ = 0.916), reflecting overall improvement within each block. However, there was no main effect of Block (*F*(1,29) = 0.44, *p* = 0.51, *η*_*p*_^*2*^ = 0.015) or interaction between Block and Phase (*F*(1,29) = 0.43, *p* = 0.52, *η*_*p*_^*2*^ = 0.015). Separate pairwise comparisons showed no differences between A_1_ and A_2_ during the early (*t*(29) = 0.92, *p* = 0.37, 95% CI = [-1.0, 2.63]°, Cohen’s *dz* = 0.17) or late learning phases (*t*(29) = -0.002, *p* = 1.0, 95% CI = [-1.75, 1.75]°, Cohen’s *dz* = 0.0004) (Figure 1F). Thus, learning was indistinguishable across A_1_ and A_2_, confirming that the intervening exposure to rotation B eliminated behavioural evidence of prior A learning.

To further test whether any residual memory of A persisted at an individual-participant level, we computed the within-subject correlation between asymptotic performance in A_1_(late learning phase) and initial performance in A_2_ (early learning phase). We reasoned that if prior A learning provided even a small retained benefit, better final performance in A_1_ would predict smaller initial errors in A_2_. However, the correlation indicated no systematic relationship between how well participants ended A_1_ and how they began A_2_ (r = 0.016, *p* = 0.94). This absence of association reinforced the conclusion that the learning of B interfered with the original A memory. We also verified that basic kinematic parameters did not differ significantly between A_1_ and A_2_. We noted that neither peak speed (*t*(29) = -0.50, *p* = 0.62, 95% CI = [- 3.55, 2.12], Cohen’s *dz* = 0.09) nor MT (*t*(29) = -1.71, *p* = 0.098, 95% CI = [-63.61, 5.69] ms, Cohen’s *dz* = 0.31) were different between the two blocks. RT, however, was smaller by about 34 ms in A_2_ compared to A_1_ (*t*(29) = -5.0, *p* < 0.001, 95% CI = [-48.26, -20.21] ms, Cohen’s *dz* = 0.91), likely reflecting practice-related improvements in movement preparation (Figure 1G-I). Taken together, these results replicate the classic A-B-A interference effect in visuomotor adaptation (Caithness et al., 2004; Krakauer et al., 2005; Kumar et al., 2018).

### Explicit learning of B did not produce interference

Having confirmed in experiment 1 that exposure to an opposing perturbation robustly interferes with prior learning, we next asked which learning mechanism gives rise to this interference. Specifically, in experiment 2 (Figure 2A) we tested whether interference would occur if rotation B was compensated via deliberate, strategy use. In this experiment, a new set of participants adapted first to rotation A (A_1_), followed by exposure to the opposite rotation (B). However, unlike experiment 1, the B learning block was designed to drive explicit strategic error compensation through a delayed visual feedback paradigm. After a 24-hour delay, participants were retested on A (A_2_).

**Figure 2.**
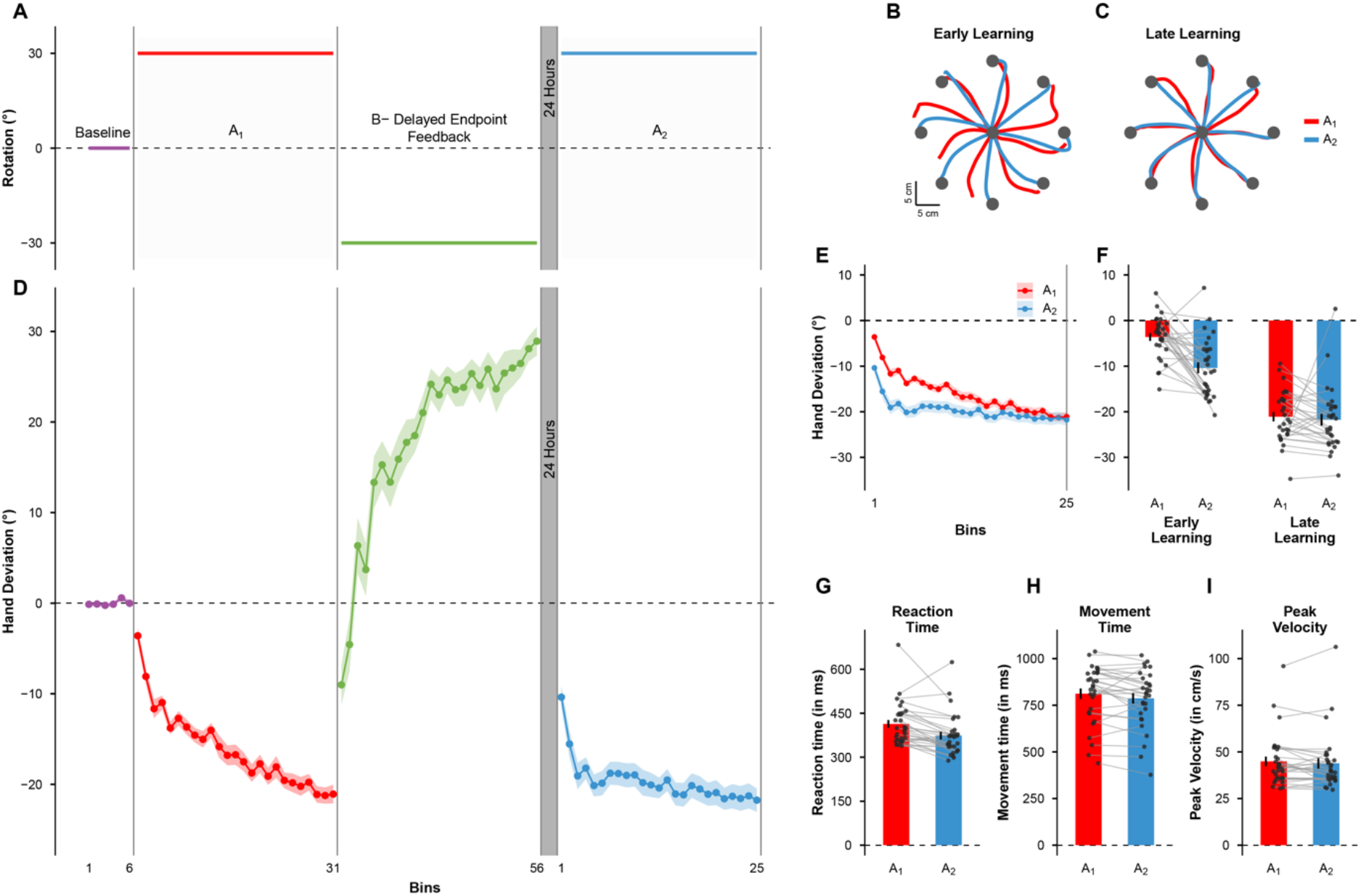
Explicit learning of B did not produce interference. **(A)** Task progression in experiment 2. Here rotation B was learned under conditions of delayed endpoint feedback. **(B)** Example cursor trajectories from a representative participant during the early phase of adaptation to A_1_(red) and A_2_ (blue). **(C)** Example cursor trajectories from the same participant during the late phase of learning in A_1_ (red) and A_2_ (blue). **(D)** Group-averaged hand deviation across bins (baseline: bins 1-6; A_1_: bins 7-31; B: bins 32-56; A_2_: bins 1-25 following the 24-hour delay). Shaded bands are SEM. **(E)** Group-averaged hand deviation curves for A_1_ (red) and A_2_ (blue) overlaid together. **(F)** Mean hand deviation during the early (first bin) and late (last bin) phases of A_1_ and A_2_. **(G)** Mean reaction time, **(H)** movement time, and **(I)** peak velocity during A_1_ and A_2_. Dots represent individual participants; bars indicate group means. Error bars represent SEM.

During A_1_, participants showed the typical pattern of gradual error reduction and trajectory straightening as they adapted to the 30° clockwise rotation (compare red trajectories of Figures 2B and 2C). Crucially, when participants were retested on A 24 hours later, the trajectories were already closer to the target direction during early learning (Figure 2C, blue). That is, subjects exhibited markedly smaller errors during the early phase of A_2_ and performance improved more rapidly, indicating that the original A memory had been preserved despite B learning. Group-averaged hand deviations confirmed this overall pattern (Figure 2D), and critically, revealed that early A_2_ movements were already biased in the appropriate compensatory direction (Figure 2E), and therefore, had smaller initial errors.

Before proceeding to statistically compare performance during A_1_ and A_2_ at the group level, we examined whether B learning indeed showed hallmarks of explicit strategy use. RT during B learning was significantly higher than during A_1_ (*t*(29) = 15.23, *p* < 0.001, 95% CI = [220.29, 288.62] ms, Cohen’s *dz* = 2.78), reflecting additional time spent in planning and implementing strategies. In contrast, MT was significantly smaller (*t*(29) = 6.58, *p* < 0.001, 95% CI = [116.39, 221.28] ms, Cohen’s *dz* = 1.20) because the absence of online feedback prevented participants from making time-consuming corrections to guide the cursor toward the target. Moreover, participants showed near-complete compensation when adapting to B, with the final hand deviation approaching 30° (mean = 28.92°, SD = 8.65°, Figure 2D, green), a feature of error compensation via explicit strategy use (Dawidowicz et al., 2022; McDougle et al., 2015). Together, these findings confirmed that B learning in the current experiment was achieved through deliberative, strategic mechanisms.

Next, to probe whether B learning interfered with the previously acquired memory of A, we compared the performance between the A_1_ and A_2_ blocks. A Block (A_1_, A_2_) x Learning Phase (early, late) repeated-measures ANOVA on hand deviation revealed a significant interaction (*F*(1,29) = 13.92, *p* < 0.001, *η*_*p*_^*2*^ = 0.324). Post-hoc tests revealed substantially larger hand deviations (smaller errors) during the early learning phase of A_2_ relative to A_1_ (*t*(29) = 5.09, *p* < 0.001, 95% CI = [4.05, 9.50]°, Cohen’s *dz* = 0.93); however, hand deviations at the end of A_1_ and A_2_ learning were not different (*t*(29) = 0.51, *p* = 0.62, 95% CI = [-2.03, 3.37]°, Cohen’s *dz* = 0.09, Figure 2F).

We also examined the relationship between asymptotic A_1_ performance and initial A_2_ hand deviations across participants. We found that this relationship was not statistically significant (*r* = 0.303, *p* = 0.103), indicating no reliable association between how well participants learned at the end of A_1_ and how they began A_2_. Finally, we compared MT and peak speed across the A_1_ and A_2_ blocks, but found no differences in either (MT: (*t*(29) = -2.01, *p* = 0.054, 95% CI = [- 49.97, 0.45] ms, Cohen’s *dz* = 0.37; peak speed: (*t*(29) = -1.06, *p* = 0.30, 95% CI = [-3.07, 0.97], Cohen’s *dz* = 0.19). As in experiment 1, RT was smaller during A_2_ than A_1_ (*t*(29) = -4.24, *p* < 0.001, 95% CI = [-59.05, -20.63] ms, Cohen’s *dz* = 0.78), reflecting increased efficiency in response initiation with practice, rather than alterations in movement execution (Figure 2G-I).

Collectively, these results indicated that explicit learning of B did not produce measurable interference with a previously acquired A memory. The original motor memory remained intact, allowing for its recall upon relearning.

### Implicit learning of B produced robust interference

Experiments 1 and 2 together established a key behavioural dissociation: interference is observed when new learning engages both implicit and explicit mechanisms (experiment 1), but not when it relies on explicit, strategic processes alone (experiment 2). This motivated us to probe whether it is implicit recalibration that is the driver of interference. To do so, we designed experiment 3, in which implicit mechanisms were selectively engaged during the acquisition of B (Figure 3A). Participants once again adapted to A first (A_1_), followed by exposure to an opposite perturbation (B). However, unlike in experiment 2, where delayed feedback encouraged explicit strategy use, here, B learning was implemented using an error-clamp paradigm designed to isolate implicit recalibration (Morehead et al., 2017).

**Figure 3.**
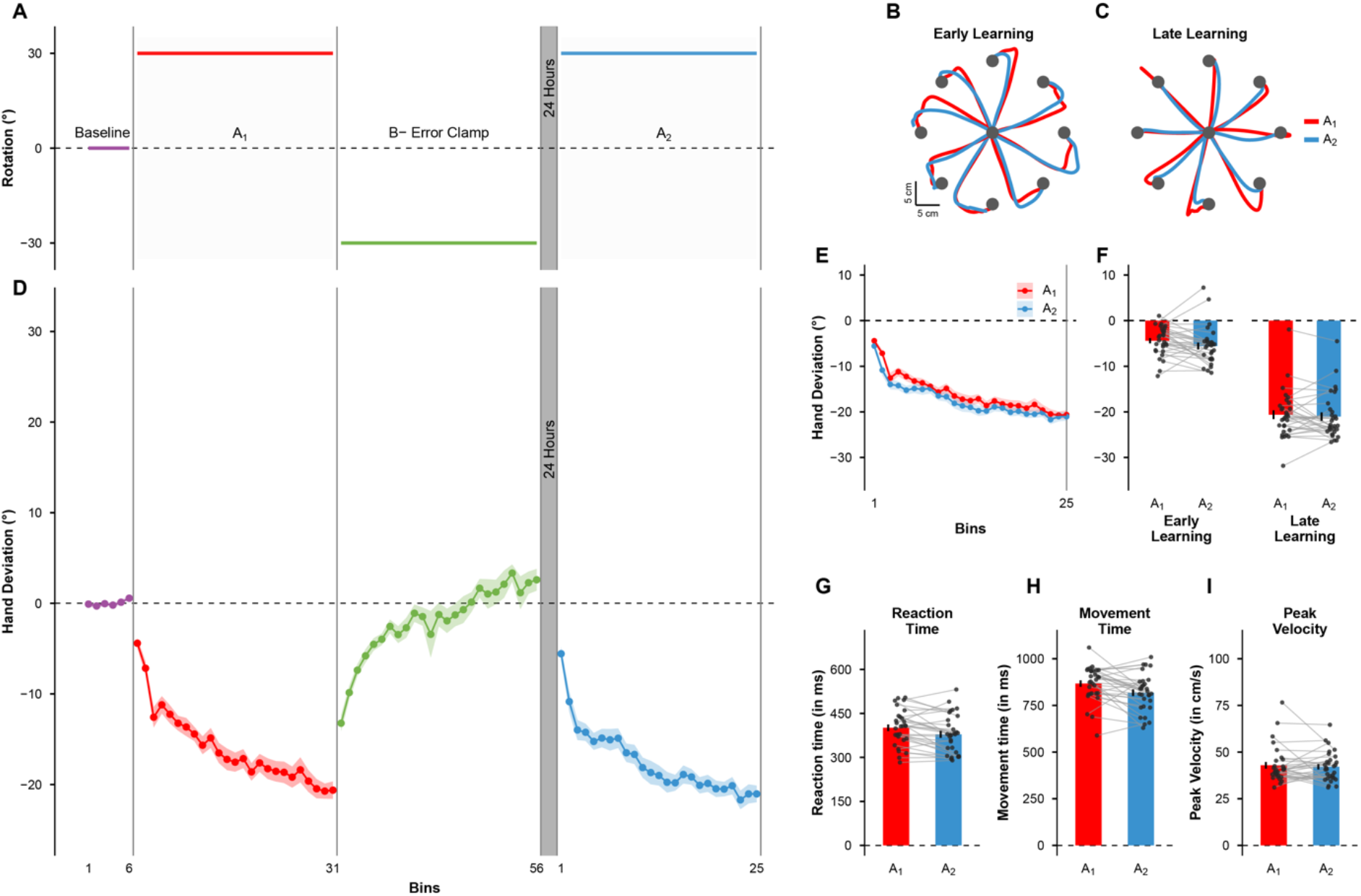
Implicit learning of B produced robust interference. **(A)** Task progression in experiment 3. Here rotation B was learned under error-clamp feedback. **(B)** Example cursor trajectories from a representative participant during the early phase of adaptation to A_1_(red) and A_2_ (blue). **(C)** Example cursor trajectories from the same participant during the late phase of learning in A_1_ (red) and A_2_ (blue). **(D)** Group-averaged hand deviation across bins (baseline: bins 1-6; A_1_: bins 7-31; B: bins 32-56; A_2_: bins 1-25 following the 24-hour delay). Shaded bands are SEM. **(E)** Group-averaged hand deviation curves for A_1_(red) and A_2_ (blue) overlaid together. **(F)** Mean hand deviation during the early (first bin) and late (last bin) phases of A_1_ and A_2_. **(G)** Mean reaction time, **(H)** movement time, and **(I)** peak velocity during A_1_ and A_2_. Dots represent individual participants; bars indicate group means. Error bars represent SEM.

For our representative participant from this experiment, trajectories were curved upon initial exposure to A_1_ (Figure 3B, red), but gradually straightened as the participant adapted (Figure 3C, red). Critically, when the same participant was retested on A after 24 hours, their hand trajectories at the start of A_2_ closely resembled those from the beginning of A_1_ (Figure 3B, blue). That is, the participant relearned A as if encountering it for the first time. At the group level, similar patterns were observed in hand deviation (Figure 3D). During A_1_, participants showed the typical pattern of gradual increase in hand deviation away from the target to compensate for the rotation. During B learning, hand direction drifted opposite to the clamped cursor. In contrast to the near-complete compensation seen when B was explicitly learned in experiment 2, the change in hand angle over the entire B learning block was only about 16° (meanΔ= 15.83°, SD = 8.22°), comparable to previously reported magnitudes of implicit adaptation (Morehead et al., 2017). Further, visual inspection of the learning curves revealed that adaptation to B progressed more slowly in this experiment than experiment 2, characteristic of the operation of an implicit process. After the 24-hr delay, performance during A_2_ closely resembled naïve learning in A_1_, with large errors occurring initially and a gradual return toward asymptote later (Figure 3E).

To test whether this implicit B learning interfered with the previously acquired A memory, we compared performance between the A_1_ and A_2_ blocks, as before. A Block (A_1_, A_2_) x Learning Phase (early, late) repeated-measures ANOVA on hand deviation revealed no significant interaction (*F*(1,29) = 0.62, *p* = 0.44, *η*_*p*_^*2*^ = 0.021) or main effect of block (*F*(1,29) = 2.76, *p* = 0.11, *η*_*p*_^*2*^ = 0.087). Only a main effect of Learning Phase was observed (*F*(1,29) = 420.69, *p* < 0.001, *η*_*p*_^*2*^ = 0.936), reflecting learning within each block. Separate pairwise comparisons confirmed that performance did not differ between A_1_ and A_2_ during either the early (*t*(29) = 1.72, *p* = 0.096, 95% CI = [-0.22, 2.53]°, Cohen’s *dz* = 0.31) or the late learning phase (*t*(29) = 0.62, *p* = 0.54, 95% CI = [-0.95, 1.77]°, Cohen’s *dz* = 0.11) (Figure 3F). This result indicated that performance during A_2_ closely mirrored naïve learning, demonstrating robust interference from the implicit learning of B.

To further examine whether any residual trace of A persisted across individuals, we computed the correlation between asymptotic A_1_ performance and early A_2_ hand deviations. Although the correlation was initially significant, inspection of the data revealed that this effect was driven by a single extreme observation with disproportionate leverage on the regression (Cook’s D = 2.164). After excluding this outlier, the correlation was no longer significant (r = 0.26, *p* = 0.173). Consistent with this conclusion, a Spearman rank correlation (with the outlier included) was also non-significant (ρ = 0.33, *p* = 0.073). Together, these analyses indicated that there was no reliable evidence for retention of A following implicit B learning, reinforcing the conclusion that interference reflects a loss of behavioural accessibility of the original memory. Finally, we compared movement-related parameters including MT, peak speed, and RT between A_1_ and A_2_. Peak speed (*t*(29) = -0.51, *p* = 0.61, 95% CI = [-3.87, 2.33]°, Cohen’s *dz* = 0.09) did not differ between blocks, but MT (*t*(29) = -2.83, *p* = 0.008, 95% CI = [-84.13, - 13.51] ms, Cohen’s *dz* = 0.52) and RT (*t*(29) = 3.03, *p* = 0.005, 95% CI = [7.23, 37.25] ms, Cohen’s *dz* = 0.55) were somewhat shorter during A_2_ (Figure 3G-I).

Collectively, these findings indicate that implicit learning of B reinstates the loss of behavioural expression of the previous A memory. The contrast between the absence of interference in experiment 2 and its reappearance here supports the view that implicit recalibration, not explicit strategy use, is the principal source of motor interference.

### Disrupting parietal processing before implicit B learning attenuated interference

Having established behaviourally in experiments 1-3 that interference arises when new learning engages implicit recalibration, we next asked whether this interference could be causally reduced by disrupting the PPC. We reasoned that if PPC-dependent processes supporting implicit recalibration drive interference, then transiently reducing PPC excitability prior to implicit learning of B should attenuate interference magnitude.

A new group of participants first adapted to a 30° CW rotation (A_1_), identical to earlier experiments. Before the subsequent 30° CCW B learning, participants received 20 minutes of cathodal hd-tDCS over left PPC. They then learned B under error-clamp conditions as in experiment 3, ensuring that learning was driven solely by implicit recalibration. After 24 hours, participants were retested on rotation A in session A_2_ (Figure 4A). All subjects performed the task only with their right arm since experiments 1-3 had shown no effect of arm used.

**Figure 4.**
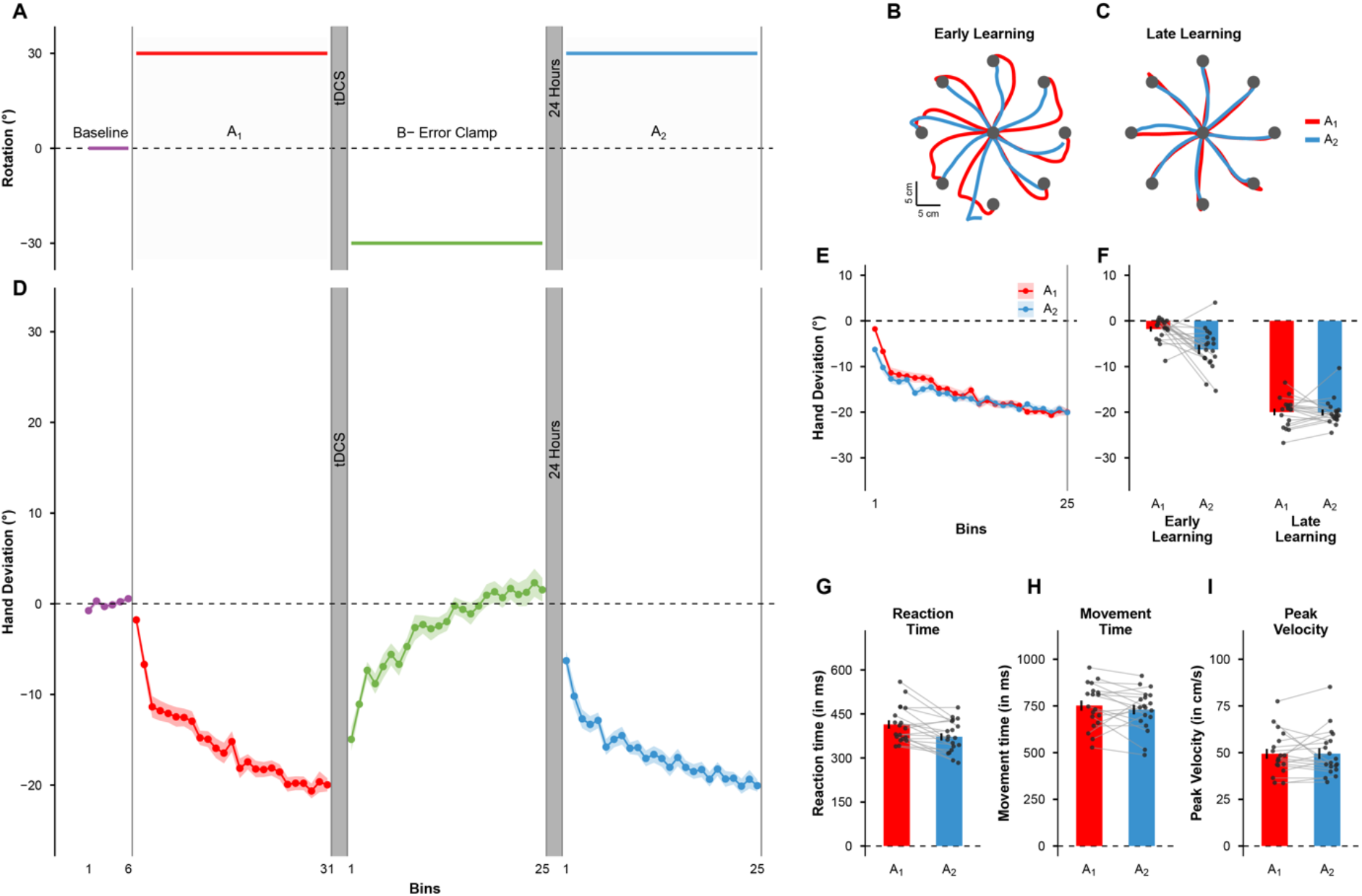
Disrupting parietal processing before implicit B learning attenuated interference. **(A)** Task progression in experiment 4. As in experiment 3, rotation B was learned under error-clamp feedback. However, cathodal hd-tDCS (2 mA, 20 minutes) was delivered over left PPC prior to B learning. **(B)** Example cursor trajectories from a representative participant during the early phase of adaptation to A_1_(red) and A_2_ (blue). **(C)** Example cursor trajectories from the same participant during the late phase of learning in A_1_(red) and A_2_ (blue). **(D)** Group-averaged hand deviation across bins (baseline: bins 1-6; A_1_: bins 7-31; B: bins 32-56; A_2_: bins 1-25 following the 24-hour delay). Shaded bands are SEM. **(E)** Group-averaged hand deviation curves for A_1_ (red) and A_2_ (blue) overlaid together. **(F)** Mean hand deviation during the early (first bin) and late (last bin) phases of A_1_ and A_2_. **(G)** Mean reaction time, **(H)** movement time, and **(I)** peak velocity during A_1_ and A_2_. Dots represent individual participants; bars indicate group means. Error bars represent SEM.

During A_1_ learning, cursor trajectories showed the classic curvature initially (Figure 4B, red), but gradually straightened as participants adapted to the rotation (Figure 4C, red). When retested on A after 24 hours, participants exhibited partial retention of prior A learning; their initial trajectories were closer to the target direction than during naïve A learning, indicating that interference was attenuated (Figure 4B, blue); trajectories also became straighter with practice (Figure 4C, blue). This pattern was consistent across participants (Figure 4D). During A_1_, the typical adaptation curve was observed, with the hand deviating away from the target to compensate for the cursor rotation. During implicit B learning, participants showed the slow, monotonic drift characteristic of recalibration, with a net change in hand angle of about 17° (meanΔ= 16.49°, SD = 6.86°), almost matching the change seen in experiment 3 (meanΔ= 15.83°, SD = 8.22°), a result confirmed statistically as well (*t*(47) = −0.29, *p* = 0.77, 95% CI = [−3.90°, 5.22°], Cohen’s *d* = 0.09). This suggested that hd-tDCS did not substantially disrupt the capacity for implicit adaptation, at least at the group level. Critically, during the early phase of A_2_, the hand direction was biased in the direction that would compensate for the rotation, consistent with reduced interference (Figure 4E).

To statistically test whether PPC stimulation resulted in interference, we compared performance via a Block (A_1_, A_2_) x Learning Phase (early, late) ANOVA on hand deviation. This analysis revealed a significant interaction effect (*F*(1,18) = 10.74, *p* < 0.001, *η*_*p*_^*2*^ = 0.374). Post-hoc tests confirmed that the early-phase A_2_ hand deviations were significantly greater compared to A_1_ (*t*(18) = 3.61, *p* = 0.002, 95% CI = [1.88, 7.11]°, Cohen’s *dz* = 0.83), whereas late-phase hand deviations did not differ (*t*(18) = 0.11, *p* = 0.91, 95% CI = [-1.59, 1.77]°, Cohen’s *dz* = 0.025) (Figure 4F). Thus, the larger deviations (smaller errors) during early A_2_ learning indicated partial retention of the A memory despite the intervening implicit B learning. For context, the average magnitude of change in hand deviation from A_1_ to A_2_ when B was implicitly learned following PPC stimulation was 4.50° (SD = 5.43°), while it was 6.78° (SD = 7.29°) when B was explicitly learned (experiment 2). This difference was not statistically significant (*t*(47) = 1.17, *p* = 0.25, 95% CI = [-6.20, 1.64]°, Cohen’s *dz* = 0.34).

We further assessed whether individual differences in A_1_ learning predicted recovery during A_2_. We found no significant relationship between asymptotic performance in A_1_ and initial performance in A_2_ (*r* = 0.20, *p* = 0.40), indicating that the retention during A_2_ was not systematically associated with how well participants adapted to A initially. Finally, MT (*t*(18) = -1.01, *p* = 0.33, 95% CI = [-66.21, 23.24] ms, Cohen’s *dz* = 0.23) and peak speed (*t*(18) = 0.03, *p* = 0.98, 95% CI = [-4.63, 4.75], Cohen’s *dz* = 0.01) did not differ significantly between A_1_ and A_2_, but as seen in our other experiments, RT (*t*(18) = 3.71, *p* = 0.002, 95% CI = [- 65.83,-18.22] ms, Cohen’s *dz* = 0.85) was smaller during A_2_ compared to A_1_(Figure 4G-I). These results indicated that stimulation did not alter movement execution, and that the observed reaction time reduction was likely due to changes in movement preparation rather than nonspecific effects of stimulation on movement execution. Collectively, these findings demonstrate that cathodal hd-tDCS over PPC delivered prior to implicit B learning is associated with a reliable attenuation of interference with the previously acquired motor memory (A_1_).

### Summary

Across our experiments, we found that motor interference depends primarily on implicit recalibration, not explicit strategy use. When adaptation engages automatic, error-sensitive recalibration mechanisms, existing sensorimotor mappings become less behaviourally accessible. In contrast, when deliberate strategies are recruited for compensating the perturbation-induced error, prior memories remain retrievable. Moreover, reducing PPC excitability attenuates this interference, implicating the PPC as a key neural substrate through which implicit recalibration disrupts previously learned motor mappings.

## DISCUSSION

Interference is a hallmark of motor memory, yet the specific mechanism responsible for interference has remained elusive. While it is well established that motor adaptation recruits both explicit strategies and implicit recalibration, recent work has largely focused on which of these processes is affected by interference. Our study was designed to invert this question to ask which process might *cause* interference. Our findings reveal a dissociation: interference is not simply an inevitable consequence of acquiring competing memories in succession, but a specific consequence of implicit recalibration. We show that when a competing task is learned via explicit strategies, the original memory remains protected. Conversely, when the competing task is implicitly learned, the original memory is no longer robustly expressed. Notably, interference depends on the nature of the learning process engaged during the intervening task, rather than on the magnitude of the competing adaptation itself.

The observed dissociation implies a fundamental difference in how explicit and implicit systems represent conflicting motor memories. Explicit strategies appear to function as context-dependent rules (e.g., “aim to the left”) that could be stored in working or declarative memory and retrieved based on task demands (Christou et al., 2016; Georgopoulos & Massey, 1987; McDougle & Taylor, 2019; Shepard & Metzler, 1971; Wang & Sainburg, 2003). In experiment 2, the strategy used for compensating the error introduced by rotation B did not overwrite the rule used when adapting to rotation A. Rather, the system likely switched contexts, preserving the A-rule for later retrieval. In contrast, implicit recalibration appears to be state-dependent and obligatory (Mazzoni & Krakauer, 2006; Oza et al., 2024; Schaefer et al., 2012; Shingane et al., 2025). Because implicit learning updates an internal model of the limb and its interaction with the environment, it may lack the flexibility required to maintain multiple, conflicting maps in parallel. Consequently, implicit adaptation to B may modify shared parameters of the sensorimotor map, rendering previously established mappings less accessible.

However, an alternative interpretation could be grounded in the multiple-model frameworks, such as the MOSAIC architecture (Haruno et al., 2001; Wolpert & Kawato, 1998) or recent Bayesian theories of contextual inference (Heald et al., 2021). In these models, the motor system stores a repertoire of context-specific internal models, and interference arises not from the overwriting of the A-model, but from a failure to retrieve it. If implicit learning of B merely suppressed the expression of prior A learning, one would predict spontaneous recovery or rapid switching given the appropriate cues. However, past work on implicit adaptation shows that it is characteristically rigid and insensitive to the very contextual cues (e.g., colour, verbal instruction) that facilitate such switching in explicit strategies (Forano et al., 2021; Mazzoni & Krakauer, 2006; Morehead et al., 2017; Oza et al., 2024; Wilterson & Taylor, 2021). This lack of contextual flexibility suggests that, unlike the modular architecture proposed by frameworks such as MOSAIC, implicit adaptation may operate on a more globally shared state estimate, rather than context-segregated representations. Within this framework then, the updates required to accommodate rotation B overwrite the parameters established for rotation A, arguing against a pure retrieval deficit explanation for the present data.

Our findings help resolve the ambiguity regarding standard interference effects, such as those replicated in experiment 1. In typical adaptation tasks, explicit and implicit processes operate in parallel. Our results suggest that the interference observed in such canonical A-B-A paradigms is driven almost entirely by the implicit process. Even if a participant retains a partial explicit strategy for the first rotation, the underlying implicit sensorimotor map could be overwritten by the intervening exposure to the second rotation, pulling the overall motor output away from the targets during retest, masking any potential savings from explicit recall. This suggests that memory stability is limited by the properties of the implicit learning system, in this case, its overwritable nature. Interestingly, prior work that has isolated implicit learning has shown that it tends to saturate at a characteristic magnitude, often well below full compensation for large perturbations, suggesting bounds on how much the underlying sensorimotor map can be updated (Albert et al., 2021; Kim et al., 2018). These known constraints raise a natural question about whether the strength of interference might depend on the portion of learning subserved by implicit recalibration. If interference reflects the degree to which implicit updates overwrite an existing map, then perturbations that fall largely within this system’s functional range may produce more complete disruption, whereas those that exceed it may yield only partial interference. Addressing this possibility will require future experiments that systematically vary rotation magnitude or error structure to determine how interference scales with the capacity of the implicit learning system.

The interference we observe here bears a conceptual resemblance to the “catastrophic interference” problem described in connectionist network models (McCloskey & Cohen, 1989; Ratcliff, 1990). In these models, sequential learning of distinct tasks involves re-optimizing shared connection weights, which can drastically degrade performance on previous tasks, reflecting a classic stability-plasticity trade-off where high adaptability compromises memory retention. In sensorimotor adaptation, implicit recalibration appears to correspond to a similar mechanism; by updating a shared internal representation, a new perturbation can overwrite a prior mapping, especially when the learning context offers no reliable cues to segregate memories. Our findings therefore place implicit motor adaptation in the same conceptual space as distributed, weight-based learning systems, highlighting the cost of plasticity when sequential tasks share a representational substrate.

That said, contemporary theories of motor memory suggest that the brain may support a more flexible, context-dependent architecture rather than simple weight updating. For instance, Heald et al. (2021) have proposed that the central computation governing motor learning is not simple weight updating, but contextual inference, a process that dynamically chooses whether to create a new memory, update an existing one, or express a previously stored mapping depending on inferred environmental context. Under this view, the motor system can maintain a repertoire of distinct internal models, each tagged to its context, allowing multiple conflicting mappings (e.g. different rotations, dynamics) to coexist without destructive interference (Hirashima & Nozaki, 2012; Nozaki et al., 2016; Sheahan et al., 2016). Our data suggest a boundary condition for this architecture. When learning relies on implicit recalibration, there appears to be limited context-based segregation, potentially leading to overwriting and interference (Avraham & Ivry, 2025). This premise implies that the protective effect of context may require explicit cues or strategic processes. In the absence of such cues, and even when adaptation engages both explicit and implicit processes (as in a number of classic studies as well as experiment 1 here), context inference may default to updating a single map. However, direct tests of contextual reinstatement or spontaneous recovery following implicit-only learning would be required to dissociate between retrieval-based and parameter-updating accounts.

More broadly, this pattern shares a conceptual resemblance with observations in declarative and episodic memory systems, where overlapping memories are often protected through mechanisms like contextual binding or pattern separation (Yassa & Stark, 2011; Yonelinas et al., 2019). In declarative memory, interference between similar events typically emerges not via erasure of prior traces but through retrieval competition or confusion when cues overlap (Anderson et al., 1994). Analogously, explicit (strategy-based) learning in motor tasks might function like a context-tagged representation. By encoding a high-level rule or cognitive strategy rather than directly overwriting the internal representation, strategy use might allow retention of prior mappings despite new learning (Forano et al., 2021). Thus, explicit strategies may play a role akin to context differentiation, enabling multiple motor memories to coexist, just as episodic memory uses context binding to preserve distinct events.

A large body of behavioural and neurophysiological data implicates a distributed cortico-cerebellar network in the kind of interference effects we see here. While it is well-established that the cerebellum drives implicit adaptation likely by computing sensory prediction errors and initiating trial-by-trial updates (Galea et al., 2011; Martin et al., 1996; Tseng et al., 2007), the posterior parietal cortex (PPC) has been proposed as a key cortical locus where these updates are integrated into more abstract representations required for motor planning (Clower et al., 1996; Kumar et al., 2020; Mutha et al., 2011; Tanaka et al., 2009). Within this framework, interference arises when new learning is successfully integrated into this shared state representation. The results of experiment 4 provide causal support for this idea; disrupting PPC processing did not abolish adaptation per se, but likely reduced the extent to which new learning was incorporated into the underlying representation. This, in turn, reduced the degree to which B learning interfered with the prior A memory.

We acknowledge the absence of a specific “sham” stimulation group in experiment 4. Although the absence of a sham condition limits causal inference, the interference observed in the “no stimulation” cohorts of experiments 1 and 3 provides a consistent benchmark against which the findings of the PPC stimulation can be interpreted. The deviation from this “standard” behaviour observed in the PPC group therefore likely represents a meaningful physiological interaction. We also consider it unlikely that the effects in experiment 4 are due to non-specific effects of tDCS. Such effects would typically result in performance changes in the *current* task (adaptation to B). However, we observed no differences in B learning, quantified as the change from the first to the last learning bin; this remained similar between the PPC group (meanΔ= 16.49°) and the “baseline” implicit group (meanΔ= 15.83°). Instead, the effect was specific to the unmasking of the prior memory. It is unlikely that a non-specific effect would selectively protect a dormant memory trace without altering the active learning task. We argue, therefore, that the attenuation of interference is most parsimoniously explained by the focal disruption of the PPC.

Finally, a point must be made about a growing theoretical synthesis that links specific behavioral hallmarks of adaptation (savings, interlimb transfer, and now interference) to distinct underlying mechanisms. Recent work has increasingly argued that savings (faster relearning) is driven by the rapid recall of explicit strategies (Haith et al., 2015; Morehead et al., 2015; Sadaphal et al., 2022). On the other hand, implicit adaptation has been identified as the primary driver of interlimb transfer, reflecting updates to an effector-independent representation (Kumar et al., 2020). Our current data suggest that this same generalized quality makes the implicit system uniquely vulnerable to interference. Because the implicit system abstracts errors away from the specific context to update a generalized map, it perhaps cannot segregate conflicting memories in the way that the strategy system can. Collectively, this body of work allows us to map features of motor memory with much greater resolution: explicit processes facilitate rapid switching and savings, while implicit processes support generalization but suffer from interference.

## Acknowledgements

We thank Gaurav Panthi for invaluable help with the hd-tDCS data acquisition, and Ajay Sahu for assistance with figure preparation. This work was supported by grants from the Department of Science and Technology, Government of India, as well as grants and fellowships from the Natural Sciences and Engineering Research Council and MITACs, Government of Canada.

